# Fast Antibiotic Susceptibility Testing based on single cell growth rate measurements

**DOI:** 10.1101/071407

**Authors:** Özden Baltekin, Alexis Boucharin, Eva Tano, Dan I. Andersson, Johan Elf

## Abstract

The emergence and spread of antibiotic resistant bacteria is a global threat to human health. The problem is aggravated by unnecessary and incorrect use of broad-spectrum antibiotics. One way to provide correct treatment and to slow down the development of antibiotic resistance is to assay the susceptibility profile of the infecting bacteria before treatment is initiated and let this information guide the choice of antibiotic. However, current methods for Antibiotics Susceptibility Testing (AST) are too slow for point of care application. Here we present a fast AST, fASTest, that rapidly captures individual bacterial cells in nanofluidics channels and monitors their response to different antibiotics based on direct imaging. By averaging the growth rate over many cells, we determined the susceptibility to several antibiotics in less than 25 min even at cell densities as low as 104 CFU/mL. The short time scale, high sensitivity and high specificity make the method practically useful for guiding antibiotic treatment in, for example, urinary tract infections.

**One Sentence Summary:** Individual bacterial cells can be captured and imaged in a microfluidic device to determine how their growth rate responds to antibiotic treatment in a few minutes.

## Introduction

With the ever-increasing emergence and spread of antibiotic resistant bacteria, a key factor in correct treatment of infections is the ability to rapidly identify the antibiotic susceptibility profile of the infecting species to assure the use of an efficacious antibiotic and reduce the need for broad-spectrum drugs (1–3). Phenotypic Antibiotic Susceptibility Tests (Phenotypic ASTs) are typically based on the detection of differential bacterial growth with and without antibiotics in liquid cultures or on solid agar plates (4). In liquid tests, detection is based on the change in optical density, while the disk diffusion method is used on solid agar plates to identify inhibition zones (5). These methods are generally reliable for detecting resistance and determining the antibiotic concentration that halts bacterial growth, making them predictive of the therapeutic utility of different antibiotics. However, since it takes 1-2 days to get a reliable readout, these methods fail to guide treatment in the early, often critical, stages of infection. As a consequence, the physician is left with the difficult choice of prescribing a broad-spectrum antibiotic or risking that the first prescribed antibiotic is ineffective.

Genotypic ASTs are based on detection of a specific genetic marker (plasmids, genes or mutations) associated with resistance phenotypes by using the common genetic tools (e.g. sequence specific amplification by polymerase chain reaction, padlock probe mediated rolling circle amplification or even whole genome sequencing) (3, 6). These tests are highly sensitive and can limit the detection time to what is needed to amplify selected DNA sequences to detectable levels, but they require advance knowledge of which resistance markers to test for. If new resistance mechanisms arise, these would go undetected and result in false negatives. Furthermore, the presence of certain resistance genes/mutations does not necessarily translate into phenotypic resistance.

Unlike the genotypic ASTs, the phenotypic ASTs directly assess if the antibiotic stops bacterial growth, which is the most relevant measure for the treating physician. New phenotypic ASTs have therefore been developed in recent years to decrease the detection times. In particular, microfluidics (7, 8) have made it possible to increase the signal to background ratio in the phenotypic assays by miniaturizing the bacterial incubation chambers (9). Using microfluidic approaches, it has been possible to push the time requirement for AST to 1-3h (10–15).

Our aim was to develop the fastest possible phenotypic AST by capturing individual bacterial cells in a microfluidic device and monitoring their instantaneous growth rate response to antibiotics. Our fluidic device allows us to filter the few bacterial cells that exist in a limited sample volume and the cell-size based growth rate determination makes it possible to monitor biological responses to the antibiotics in a few minutes.

## Results

### Design and Loading of Microfluidic Chip

A simplified design of the microfluidic chip is shown in Fig.1. (More details on the design and alternative operation modes can be found in the supporting information (SI)). The device has two rows of cell traps. In each row, there are 2000 cell traps of dimension 1.25µm x 1.25µm x 50µm (Fig. 1A.) that can be loaded with bacterial cells (red) from the front channel. A constriction at the end of each trap prevents cells from passing to the back channel while allowing the media to flow around the cells. This sieve-like design, which is an extension of the Mother Machine (MM) (16), enables rapid loading and constant media flow over the cells.

**Fig. 1.**
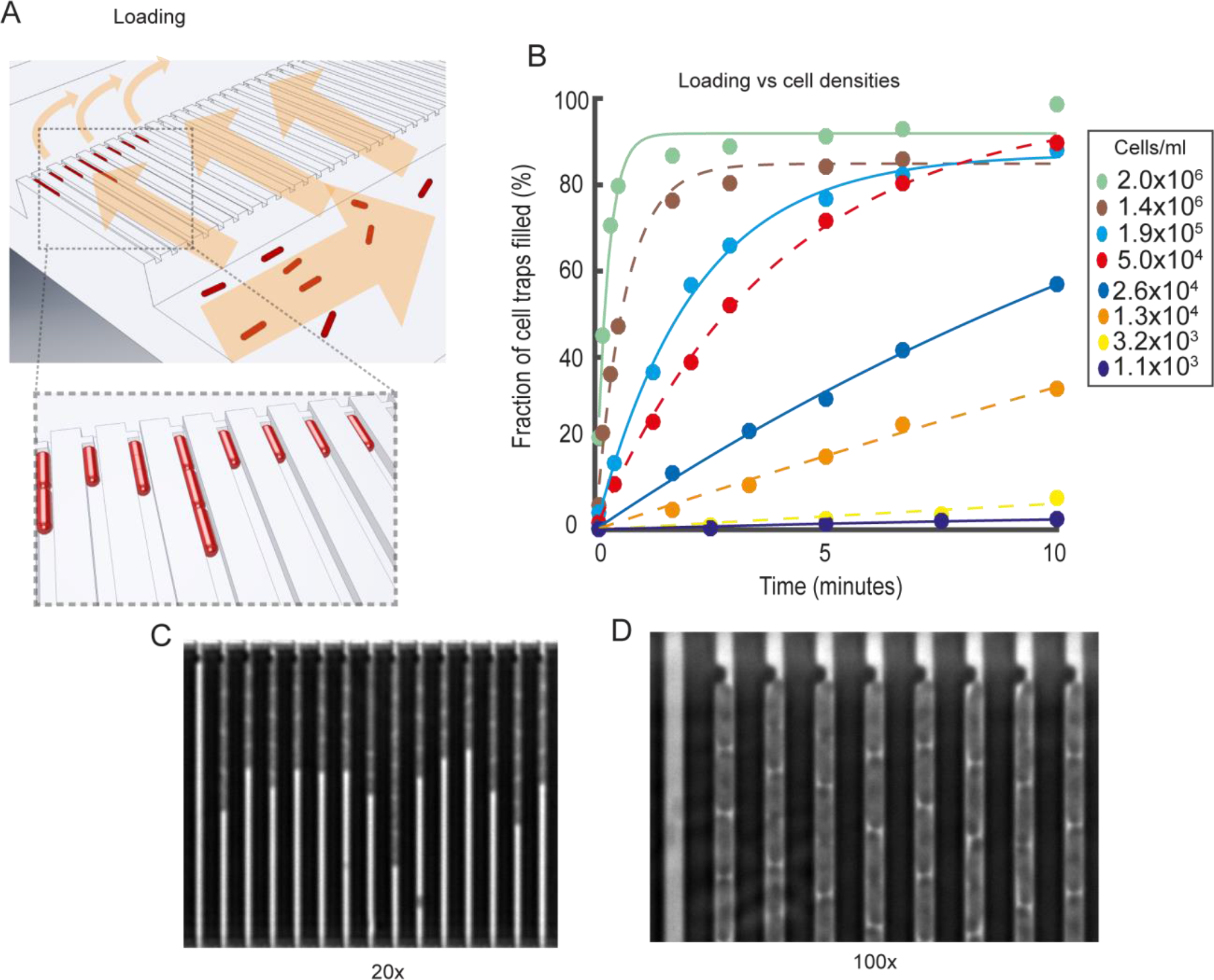
Design and operation details of the microfluidic chip used in the fASTest. (A). Cartoon illustrating the loading of rod shaped bacterial cells (red) into cell traps. Arrows indicate flow direction during loading (B) Fraction of cell traps with at least on E. coli cell at different time points. The different markers correspond to different density cell cultures (C) A phase contrast image of E. coli in the microfluidic device (darker regions) using a 20x objective. (D) A small part of a phase contrast image taken at 100x during fASTest showing the back end of the cell trap where the flow restriction region captures the cells during loading.

In order to test how rapidly dilute samples could be loaded, we monitored the fraction of traps loaded over time for different cell cultures with densities from 1100 CFU/mL to 2 × 106 CFU/mL (Fig. 1B). 80% of the cell traps were filled within 30 seconds at a cell density of 2 × 106 and even with the most dilute sample, 1100 cells/mL, 160 traps were filled in 10 min. (SI section: Results of Low Cell Density Cultures. Fig. S4).

The loading time curves (Fig. 1B) can also be used for real time bacterial density determination, for example in a urine sample. Within 30 seconds it would be possible to assess if it was a severe bacterial infection with >105cells/mL and within 10 min it would be possible to determine if there is a clinically relevant infection at all (>103cells/mL urine) (17, 18).

### Growth rate measurements

In a typical experiment, fluid control is switched from Loading to Running mode when half of the cell traps are loaded. A few seconds after the switch, loading media inflow stops and test media starts to reach the cell trap regions. The reference population row receives the antibiotic free medium and the treatment population row receives the media with the antibiotic to be tested (Fig. 2A). Following the media switch, we performed time-lapse phase contrast microscopy (20x magnification) with an automated X-Y translation stage, where each of the 4000 cell traps were imaged every 60s. (A time-lapse movie of one position during a typical experiment can be seen in SI Movie 1).

**Fig. 2.**
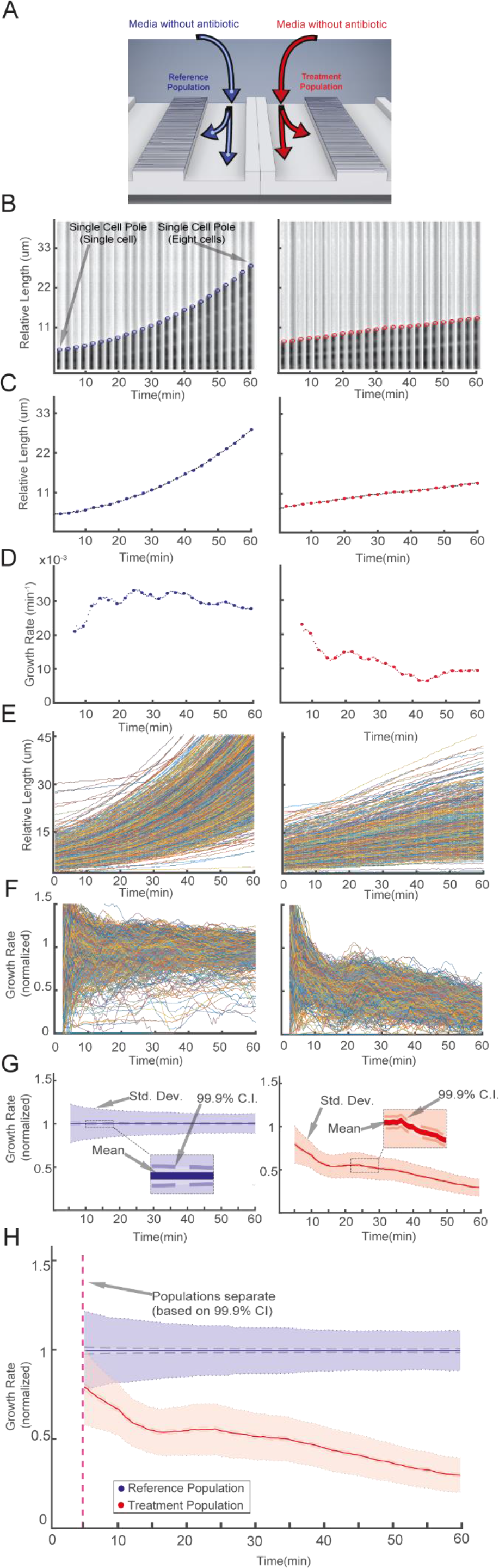
Detection of Growth Rate Effect of Antibiotic. (A) Media with or without antibiotic is supplied to the two different rows of cell traps to test the effect of the antibiotic (Ciprofloxacin,1 µg/mL) on the treatment population compared to the reference population. (B) A single cell trap from the reference population (left), and another single cell trap from the treatment population (right) are shown every 5^th^ frame (every 2.5 min). The detected front-most cell pole position is given as a blue or red circle. (C) The corresponding length change through time is shown with the blue or red dots. (D) Growth rates calculated with a sliding window of up to 10 min. (E) The length and (F) growth rate as a function of time is plotted for all the individual traps for the reference (left) and for the treatment population (right). (G) The descriptive statistics of the normalized growth rate distributions for these populations are shown as a function of time. Colored as blue for the reference population (left) and red for antibiotic treated population (right), solid lines show the mean, dark shaded region show the 99.9% CI of the mean and the light shaded region show the sample standard deviation. (H) On the overlaid figure of two population’s normalized growth rate distributions, time of separation of the treatment population from the reference population based on 99.9% CI is given as dashed magenta line.

Growth rates were first estimated for the cells in each individual cell trap. The movement of the front-most cell pole in each cell trap was monitored as a proxy for the cumulative cell length changes for the cells within that trap. The top part of the Fig. 2 (B, C and D) shows an example of how the average growth rate for the cells in an individual cell trap was calculated for the reference cells without antibiotics (left) and antibiotic treated cells (right). Fig. 2B show one cell trap every 5^th^ frame of a 120 frame (60 min) experiment and the circles (blue or red) show the detected front-most cell pole position at that frame. In Fig. 2C, the solid dots (blue or red) show the length of the cell(s) within that cell trap through the experiment. Fig. 2D shows the instantaneous growth rate estimated as the relative length increase per time in a sliding window over 10 minutes.

### Distinguishing two populations based on growth rate statistics

In the lower part of the Fig. 2 (E, F, G and H) we applied the growth rate calculation for each of the 2000 cell traps in both rows such that the Fig.2 E shows length versus time for the two-population (reference population on the left, treatment population on the right). The corresponding growth rates for individual traps are shown in Fig. 2F. In Fig. 2G, the growth rates for the individual cell traps from the both populations are averaged and normalized by the average growth rate of the reference population. The frame-wise sample standard deviation is shown in light transparent color and the standard error of the mean is used to calculate a 99.9% confidence interval for the average growth rate (darker shaded region). By combining the data from the two columns (Fig. 2H), we could detect the response of the antibiotic treatment population in this experiment based on 99.9% confidence intervals’ divergence (dashed magenta line) earlier than the first data point at ∼4min.

### Fast detection of response to antibiotic treatment

Urinary tract infection (UTIs) is one example where fast AST could improve medical practice by making it possible to prescribe an antibiotic to which the infecting bacteria are susceptible before the patient leaves the primary care unit. Using the fASTest, we determined the antibiotic response time of E. coli (MG1655) to nine different antibiotics that are used for urinary tract infections. These included three different types of Penicillins (AMP: Ampicillin, AMX-CLA: Amoxicillin-Clavulanate and MEC: Mecillinam), one Carbapenem (DOR: Doripenem), two different Fluoroquinolones (CIP: Ciprofloxacin and LEV: Levofloxacin) and three other agents (FOS: Fosfomycin, NIT: Nitrofurantoin and TMP-SMX: Trimethoprim-Sulfamethoxazole). We followed EUCAST breakpoint value recommendations for the test concentrations of each particular antibiotic and for the testing media requirements. We loaded the bacteria into the microfluidic chip and supplied growth media without antibiotic to one row (reference population) and growth media with antibiotic to the other (treatment population). With this set-up, we could detect the differential growth rate between treatment and reference populations in 3 min for CIP, LEV, MEC, NIT, TMP-SMX; in 7 min for AMX-CLA and DOR; in 9 min for FOS; in 11 min for AMP based on 99.9% confidence intervals (Fig. 3).

**Fig. 3.**
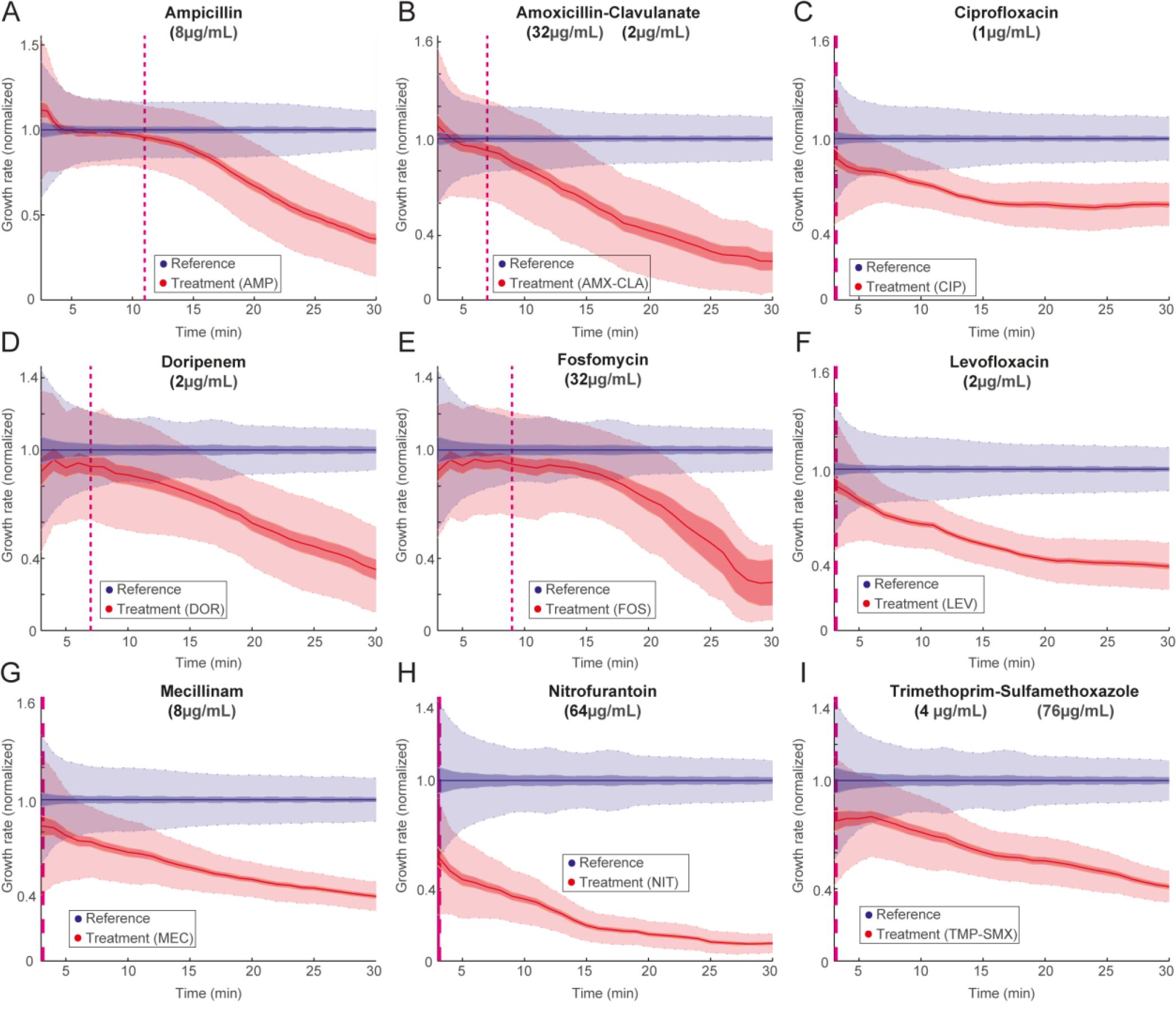
Fast detection of response to antibiotic treatment. fASTest experiments testing how fast susceptible *E. coli* cells responds to Ampicillin (A), Amoxicillin-Clavulanate (B), Ciprofloxacin (C), Doripenem (D), Fosfomycin (E), Levofloxacin (F), Mecillinam (G), Nitrofurantoin (H), Trimethoprim-Sulfamethoxazole (I).

Importantly, the response curves were very reproducible between biological replicates, as shown for CIP and AMP in the supplementary material (S.Fig.5). This implies that the differential responses to the different drugs were not due to limited time resolution in the assay but rather that the measurement was sufficiently fast to monitor the actual biological response time. These experiments were run in the standard susceptibility testing MH-media, however in the supplementary material we show that response times were similar for bacteria grown in urine (S.Fig.6). In the supplementary material, we also show that the fASTest works on *Klebsiella pneumoniae* and *Staphylococcus saprophyticus*, which are two other major pathogens causing uncomplicated UTI. The response time for *S. saprophyticus* was slower as expected due to its longer generation time (S.Fig.7).

### Fast Antibiotic Susceptibility Testing of clinical isolates

To be a practically useful AST, fASTest needs to differentiate strains with a clinically relevant spectra of resistance mutations from susceptible strains in their responses to the antibiotic. This is important since even resistant bacteria can show a growth rate reduction due to presence of an antibiotic, and one can therefore not directly conclude that a strain is susceptible just because it has impaired growth compared to untreated cells (as seen in Fig 3). For this reason, we first tested an E. coli strain that was genetically engineered to be resistant to ciprofloxacin (CipR) due to the introduction of several clinically observed mutations (gyrA1-S83L gyrA2-D87N parC- S80I) (19) for Ciprofloxacin susceptibility and compared the results with that of the susceptible wild type E. coli strain (CipS) (Fig. 4A and B). During the first minutes of the test the CipR strain grew slightly slower when treated with CIP than its untreated reference population (Fig. 4A) before recovering to the same growth rate. The CipS strain responded more strongly (Fig. 4B), which made it possible to distinguish the wild type strain from its constructed resistant counterpart in a few minutes.

**Fig. 4.**
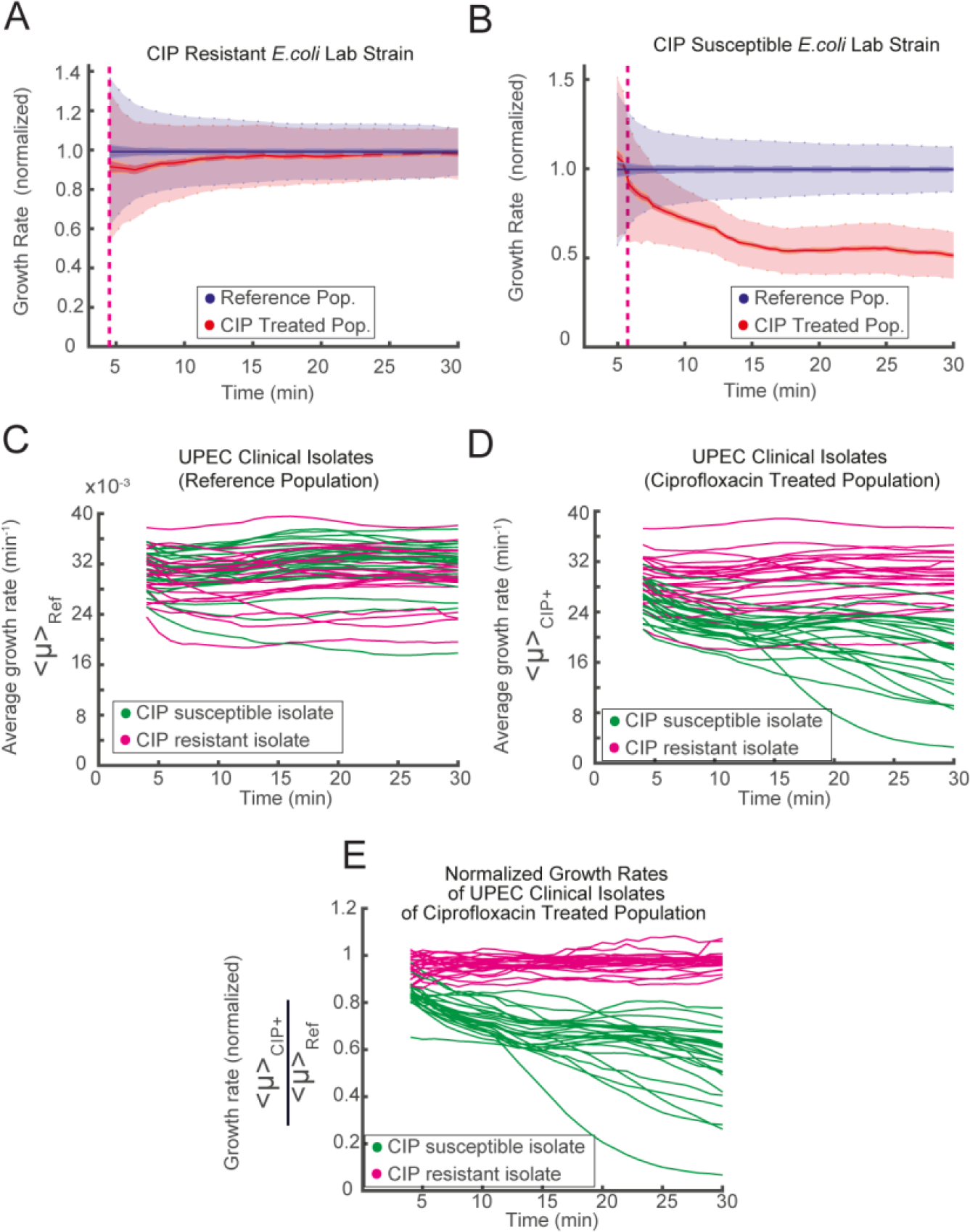
fASTest for resistant and for susceptible strains. Lab strains of Ciprofloxacin Resistant and susceptible (B) *E. coli* are tested for CIP susceptibility. (C-E) 49 Clinical Isolates of Uro-Pathogenic *E. coli* (UPEC) are tested with fASTest for CIP susceptibility; (C) Average growth rates for the reference populations (D) Average growth rates for the treatment populations. Color coding: magenta for clinically resistant and green for clinically susceptible to Ciprofloxacin. (E) Growth rate of treated populations normalized for the growth rate of the respective reference population.

Next, we tested 50 clinical isolates of uropathogenic E. coli (UPEC) strains for Ciprofloxacin susceptibility. Samples were initially collected and characterized by the Uppsala University Hospital Clinical Microbiology Laboratory using the gold standard disk diffusion test for CIP susceptibility/resistance as a part of routine for clinical diagnosis. The antibiotic susceptibility of isolates was tested by disk diffusion method recommended by the European Committee on Antimicrobial Susceptibility Testing (5). 25 CIP susceptible and 25 CIP resistant isolates were given to us. We received those samples blinded, without the patient nor the CIP susceptibility information. One isolate did not grow in the liquid cultures during the preparations and was therefore excluded from the study. The remaining 49 isolates were tested for CIP susceptibility using the fASTest. Afterward, the results were compared to the results from the hospital.

The fASTest results of 49 UPEC clinical isolates are given in Fig 4. C-E. The growth rate of the untreated reference population varied between the isolates without any obvious differences between a resistant and susceptible isolate (Fig 4.C). CIP Growth rates of treated populations did not reveal a striking distinction between the resistant and the susceptible isolates (Fig 4.D). However, when the growth rate of the CIP treated population was normalized with the growth rate of untreated reference population from the same experiment, susceptible and reference isolates grouped apart after the first 10 minutes (Fig 4.E). As can be seen in Fig 4E, all the 24 resistant strains and 25 susceptible strains were correctly grouped within 10 minutes. This complete agreement corresponds to 86.28% to 100.00% sensitivity and 85.75% to 100.00% specificity for detecting resistance.

## Discussion

This study originated in an investigation of the cell to cell variation in the bacterial cell cycle (20), which required us to develop new tools to determine growth rates very accurately for individual *E. coli* cells. The process of averaging the growth rate from several hundred individual bacteria makes it possible to detect changes in growth as fast as the biological responses to the antibiotic (S.Fig.5). For this reason, it is not possible to find a phenotypic AST based on the growth rate response that is faster than the fASTest. It is important to note that the same time resolution cannot be achieved in a bulk measurement of the growth rate since in that case the averaging is done over the cells before the readout, which means that the unavoidable readout noise will impact the growth rate much more than if it is averaged out over the cells after the readout.

The sensitivity of fASTest in terms of how many cells per ml can be detected is only dependent on how much time is spent loading the cells. For 104 CFU/mL, which is in the lower range for clinically relevant UTIs, sufficient loading can be achieved in 5min, since for fASTest it is sufficient to collect approximately 100 bacteria. However, the sensitivity will practically depend on which body fluid the sample is collected from. For examples, if there is a large number of bacterial sized contaminants that clog the traps in the chip, it will be the concentration of contaminants that determines the sensitivity.

We have here focused on bacterial species and antibiotics related to UTIs, but it is likely that the same principles would work for sepsis or meningitis, i.e. in blood or cerebrospinal fluid that normally should be devoid of bacterial sized cells. Independent of the sample, the key principle will be true: it is sufficient to measure the single cell growth of a few hundred bacteria to get very close to the theoretical time limit for monitoring the response to an antibiotic in real time.

## Materials and Methods

### The microfluidic chip

The microfluidic chip consists of a cover glass (#1.5) and a micro-molded silicon elastomer (Sylgard 184, PDMS) that are covalently bonded together. For micromolding we used the standard soft lithography techniques as described in the SI Materials and Methods.

### fASTest Protocol

All the fASTest runs described in this paper follow this common protocol: Growth of over-night culture, growth of loading culture, connection of microfluidic flow control setup to the microfluidic chip, positioning of the chip according to the camera, selection of positions to be imaged, running an imaging test to ensure the stability of the microfluidic chip, connecting the loading culture to the macrofluidic setup, loading the cells from loading culture, starting the antibiotic application and automated phase contrast microscopy.

### Bacterial Strains

The strains used were wild-type (wt) strain: DA5438 (*Escherichia coli* MG1655). Ampicillin resistant strain: DA28097 (*E. coli* del(P*lacI*_*lacIZYA*)::amp). Ciprofloxacin resistant strain: DA20859 (*E. coli gyrA1*-S83L, *gyrA2*-D87N, parC-S80I). Two other species:

DA12755 (*Klebsiella pneumoniae,* ATCC13883), DA14015 (*Staphylococcus saprophyticus*).

### Growth Media

Depending on the experiment, we used either Mueller-Hinton Broth (Sigma-Aldrich 70192- 500G) or Urine as Growth Media (GM). When indicated, the GM was supplemented with an antibiotic. In preparation of urine media, morning urine was collected, centrifuged for 30min at 4°C to get rid of the large debris by pelleting and then filtered (nitrocellulose filter, 0.2µm pore size). All media is supplemented with a surfactant (Pluronic® F-108, Sigma #542342, 0.85% (w/v) final concentration) to prevent the attachment of the bacteria to the PDMS surface.

### Culture Conditions

Over-Night Culture (ONC): Bacteria from the glycerol stocks were inoculated into 2ml Growth Media (GM) and incubated (37˚C, shaking 225rpm) for ∼16h. Loading Culture: 2.5µl ONC is diluted 1:800 to total 2ml GM and incubated (37˚C, shaking 225rpm) 120min. Growth in the chip: The chip was continuously supplied with GM and incubated in the microscope cage incubator at 37˚C before, during and after the loading of bacterial culture, and also during the test. The loading culture was connected to the fluidic setup and kept in the cage incubator at 37˚C. GM was kept outside of the cage incubator at room temperature (21˚C).

### Microfluidic flow control setup details

Flow direction and rate during the experiment were maintained by pressure driven flow. An electro-pneumatic controller from Elveflow (OB1 MkIII) regulated the air pressure applied to the closed fluidic reservoirs. Pressures, flow rates and tubing details are explained in SI Materials and Methods. The electro-pneumatic controller was programmed in MATLAB.

### Automated Phase Contrast Microscopy

We used a Nikon Ti-E inverted microscope with 20X Objective (CFI Plan Apo Lambda DM 20X or CFI S Plan Fluor ELWD ADM 20X), phase contrast module, motorized X-Y Stage, Perfect Focus System (PFS) and a CMOS camera (The Imaging Source, DMK23U274). The setup was maintained within Cage Incubator Enclosure (custom made by Okolab), where the temperature was maintained at 37°C by a temperature controller (World Precision Instruments, Airtherm-Atx). Both the microscope and the camera was controlled by an open source microscopy software (MicroManager 1.4.19). Phase contrast images were acquired by the software’s multidimensional acquisition feature through which the motorized stage move the fluidic chip to 36 different positions. Each position was imaged every 30 seconds or every 60 seconds depending on the particular experiment. Each experiment was 30 minutes, even though the imaging could be continued longer if needed to provide insights on kill dynamics.

### Image Processing

The images were processed for detection of each row in the raw image, cell traps and empty traps in each row, removing background and performing pole detection to obtain the cell pole detection in each frame of each position using an algorithm developed in MATLAB. Details of this algorithm is given in SI Materials and Methods.

### Data Analysis

For cell pole tracking we used µTrack (21). For growth rate calculation of individual cells, we applied a sliding window of data points (length) and fitted a linear function to the logarithm of them. The sliding window grows from 2minutes to 10 minutes in the beginning of the experiment and stay at 10 minutes afterwards. We filtered some data based on fixed criterion to remove misidentified particles or cells that were dead from the beginning as well as the traps that were overly filled or left empty during the loading. Details of filters applied are given in SI Materials and Methods.

## Supplementary Materials

- Materials and Methods
- Fig. S1. Chip Design and features
- Fig. S2. Mold scanning electron microscope images
- Fig. S3. Image processing steps
- Fig. S4 fASTest low cell density cultures
- Fig. S5 fASTest repeatability
- Fig. S6 fASTest Resistance detection in Mueller-Hinton and in Urine for CIP-R
- Fig. S7 fASTest using 2 other species
- Movie S1. Time lapse images of an example fASTest for WT E. coli
- Movie S2. Time lapse images of Staphylococcus saprophyticus growing in chip
- Movie S3. Time lapse images of Klebsiella pneuomoniae growing in chip

## Acknowledgments

### Funding

This work was supported by the European Research Council (J.E.), the Swedish Research Council (J.E. and D.I.A.), and the Knut and Alice Wallenberg Foundation (J. E.).

## Author contributions

Ö.B., D.I.A. and J.E. designed the research; Ö.B. performed the experiments; D.I.A. provided the bacterial strains; E.T. provided the UPEC clinical isolates; A.B. and Ö.B. contributed new analytical tools; Ö.B. analyzed the data; and Ö.B., A.B., D.I.A. and J.E. wrote the paper.

## Data and materials availability

All raw data will be made available at request for non-commercial interests. **Ethical permit:** The ethical review committee in Uppsala has no objection against this study (ref 2017/051).

## Supplementary Materials

### Materials and Methods

#### Chip Design and Features

Here we describe the features of the chip that we have designed to improve the capturing, imaging and manipulation of individual bacteria. We refer to the general chip design as xGMM, where the x refers to the number of parallelization option for a given Mother Machine (MM) chip. G refers to the Growth Condition. The multiple modes of operation are explained in the next section.

The main feature of xGMM consists of a cell trap region (Fig. S1. C), with cell traps parallel to each other and perpendicularly connected from both ends to a front and a back channel that are parallel to each other. The cell trap size can be manufactured according to the dimensions of the cells to be studied. For the *E. coli* that is studied in this paper we used 50µm long cell traps with either 1.25µm x 1.25µm cross sectional area. The pitch can also vary according to the purpose of the chip but in this study, we used chips with cell traps every 3 or 4µm.

The front and the back channels have a fluidic connection to each other only through the cell traps. In each cell trap, 3µm away from the point that is connected to the back channel, there is a constriction region of 1µm length, where the width of the cell trap is 300nm. *Escherichia coli* cells are captured as they cannot easily pass through the constriction region when they flow from the front channel to the back channel through the traps during loading.

To prevent the flow of the large particles (i.e. dust, precipitates in the growth medium) and to capture large air bubbles, we have included a filter region (Fig. S1. B) at each fluidic port. The filter region has different zones that gradually decrease the filtration size to prevent clogging. In sequence, it captures particles bigger than 160, 80, 40, 20 and 10µm at different parts of the port and prevents anything bigger than 10µm from flowing in. The distances between the different filtration zones are designed to be the size of the next filtration region thereby creating an extra filtration effect between two zones (e.g. distance between 160µm zone and 80µm zone is 80µm). This also increases the filtration capacity.

Some features of the chip are designed specifically to aid faster image acquisition or more precise image processing algorithm. These features are the empty traps, the dot-code regions, and two parallel cell trap rows located close enough to each other to fit into the same field of view in 20x magnification.

At a fixed periodicity, there are built-in empty traps, (Fig. S1. C) that have constrictions in the front end of the traps instead of the back, preventing the entry of any cell during the loading. The empty traps are used by the image processing algorithm for background subtraction (see below) which increases the contrast and makes it easier to detect cell poles.

A dot-barcode is imprinted next to each empty trap. The dot-barcode is used to identify the indices of the adjacent cell traps using a binary code consisting up to 12 dots (Fig. S1. C), 10 dots for the binary code and one top and one bottom dot to specify the positioning of the code region.

**Fig. S1.**
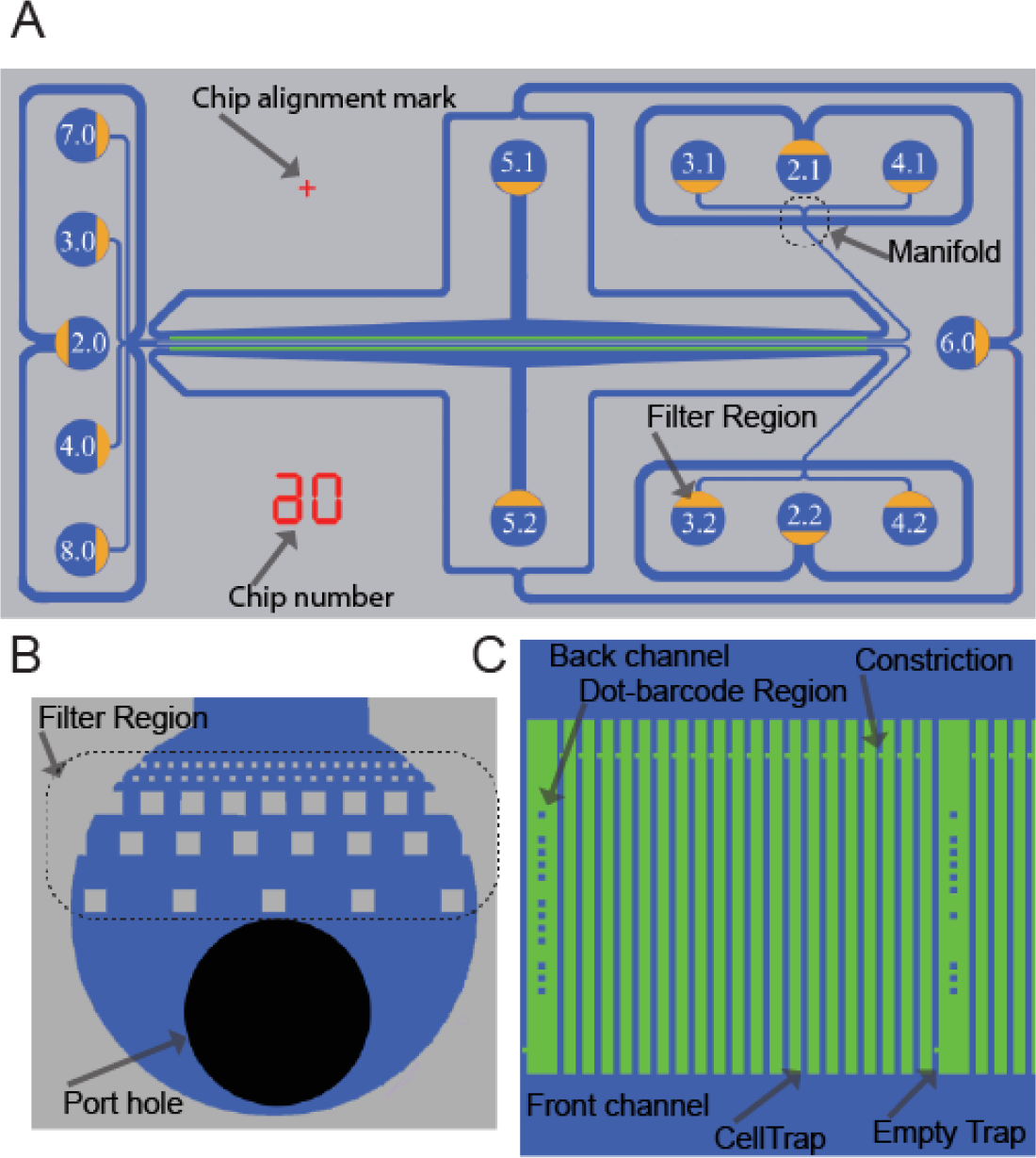
2GMM Design and features. (A) Drawing of the 2GMM design showing all the ports with designated port numbers and the orientation of filter regions (yellow) in the ports. Chip number and chip alignment marks are shown red. (B) Drawing of a port showing the port-hole (black) and the filter region. (C) Drawing of a section of a cell trap row.

The two cell trap rows of 2GMM are located 150µm apart from each other and are parallel. In the field of view of an 20X magnified phase contrast image covering ∼261 × 348µm all of these structures can fit for a ∼100 cell trap wide section of the chip from both of the rows. Thus, during the automated image acquisition, the motorized X-Y stage of the microscope needs to mainly move in X-axis, which maximizes the positions that can be imaged in a given time and thereby minimizing the frame rate and increasing the time resolution.

### Alternative Modes of Operations and Parallelization

In each 2GMM, there are two distinct cell trap regions, also mentioned as cell trap rows (or simply rows). Fluidic isolation of these rows from each other enables the delivery of two distinct fluid streams simultaneously, which permits parallel testing. If the aim is to test one strain with two different growth media condition in parallel, then the running step benefits from the fluidic isolation of the two rows and each growth media is applied to a different row. For example, for the fASTest we use the 2GMM by punching holes only in the necessary ports 2.0 (cell culture loading port), 5.1 and 5.2 (wash out ports), 2.1 and 2.2 (waste ports) and the 3.1 and 3.2 (media ports) (Fig. S1. A). Meanwhile, all other ports are not used.

### Mold Production

Molds are produced by the company NMetis (Göteborg, Sweden) according to our design. The substrates are 3-inch Si-wafers of 1000µm thickness. The mold is produced in 3 stages. First, a SiO2 layer is formed using thermal oxidation. Second, cell traps and optical alignment marks are etched in the oxide layer using combination of electron beam lithography, reactive ion etching and wet etching. Third, the lithography of flow channels, ports and filtering regions are made in 10µm thick SU8 layer. Some of the features of the 2GMM mentioned in the first section can be seen from the scanning electron microscopy (SEM) images of the 2GMM Mold (Fig. S2.)

### Micromolding

The microfluidic chip consists of two parts covalently bonded together in the lab: a cover glass (#1.5) and a micro-molded silicon elastomer (Sylgard 184, PDMS) that a covalently bonded together. For micromolding we used the standard soft lithography techniques. Polydimethylsiloxane (PDMS), Sylgard 184 (Dow Corning), base and the curing agent is mixed thoroughly at 1:10 ratio using a homogenizer (FastPrep-24. MP Biomedicals). The mixture is degassed by centrifuging at 1878g for 30sec. After pouring the mixture into the molds, it is further degassed under vacuum. Curing is done at 100°C for 1- to 15h. Demolding is followed by dicing micro-molded arrays of PDMS Chips into individual PDMS chips, punching fluidic connection ports, and cleaning the surface of PDMS chips. Cover glasses were pre-cleaned under running de-ionized water from dust and particles and then they were incubated in glass cleaning liquid (0.2% v/v Helmanex III in Mili-Q water) in a Teflon vessel with sonication for 45min and rinsed thoroughly with and stored in Mili-Q water. Before bonding, both the PDMS’ and the cover glass’ surfaces are cleaned and activated. PDMS surfaces are cleaned by Scotch Magic Tape (3M) initially and followed by an isopropyl alcohol (Sigma-Aldrich) dip and blow dried using clean pressurized air. Surface activation is done using air plasma treatment using HPT-200 Benchtop Plasma System (Henniker Plasma) at half power (100W) for 30sec. The surfaces are cleaned using a UV-Ozone Cleaner for 10min. Right after that, the surface activation is done by applying a local corona discharge from a high frequency spark generator by scanning the application probe over the surfaces for 30-60sec staying ∼within 5-10mm of the surfaces. The activated PDMS surface is placed on the activated surface of the cover glass to initiate the bonding. After initial attachment, the microfluidic chip is placed in 100°C for at least 30min.

### Macrofluidic Setup Details

The fluid reservoirs are 15ml Falcon tubes with reusable reservoir adapters (Elveflow). Tygon tubing (inner diameter (ID) 510µm, outer diameter (OD) 1524µm) (Saint-Gobain) is used to connect each reservoir to the chip (length 100cm). The tubing is connected to the chip using a custom-made metal connector (23G, 14mm long tubing bend in the middle with 90 degrees) (New England Small Tubing). During loading, both the media and the loading reservoirs are pressurized with 500 mbar and during the running, only the media reservoirs are pressurized with 100mbar. Pressure control is done using OB1-Mk3 from Elveflow. Waste ports are connected to a 10cm tubing, the other end of which is open and in level with the chip.

**Fig. S2.**
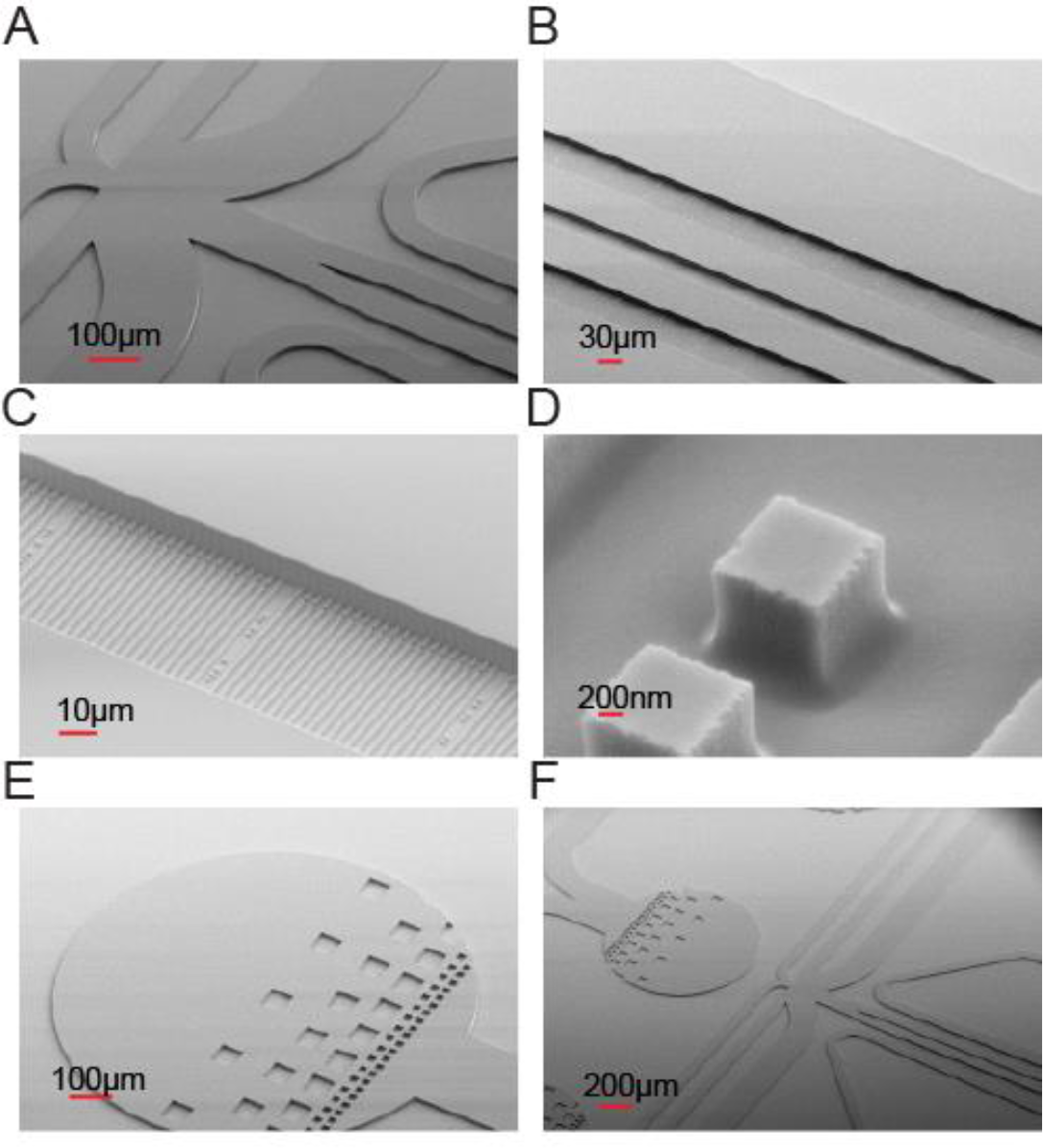
2GMM Mold scanning electron microscope images with different magnification shows some features of 2GMM on the mold such as (A and F) manifold, (B) two rows of cell traps, (C) cell traps, (D) dot-barcodes, (E and F) filter regions on the ports.

### Image Processing

For image processing, we developed an algorithm encoded in MATLAB. The algorithm has subsections with different functions as explained below.

#### Note on representation of images in MATLAB

A two-dimensional image with pixels is represented in MATLAB as a 2-dimensional matrix of elements arrayed in columns and rows. Location of each pixel on the x-axis of the image is represented in the column number of the element, and on the y-axis of the image is represented in the row number of the element. Each element’s value represents the (16-bit) gray value of the corresponding pixel.

#### Region of Interest (ROI) detection

The bottom and top rows need to be detected and cropped. The known physical distance between the two rows means that only one row needs to be precisely detected, here the top row (higher in y value). A first estimate of the location of the top row is obtained by detecting the bright horizontal lines by summing the pixel values along the x-axis of each frame to obtain the intensity profile along the y-axis of the image region (Fig. S3. A). This profile is then smoothed to flatten noise and, the two top (highest y value) lines provide an estimate of the location of the top row. The more precise estimate of the location is obtained by detecting the top dot of the bar code located between the cell traps. The dots are located using a radial symmetry dot detection algorithm *(33)*. The median location of all top dots provides a precise location of the row in the image region (Fig. S3. C).

#### Cell trap detection

Once the rows are cropped, the cell traps are detected. For the first frame, a first estimate of the locations of the cell traps is computed by summing the pixel values of the image along the y-axis to get the intensity profile along the x axis, and detecting the peaks in the intensity profile region (Fig. S3. B). As the cell traps are brighter than the rest of the image, on the intensity profile, they appear as peaks of high intensity. For all subsequent frames, the precise location from the previous frame is used as a first estimate. Using this first estimate based on the previous frame, the precise locations of the cell traps are obtained in the current frame by computing the cross correlation between each cell trap region and a cell trap mask, a binary image depicting a rectangle the size of a cell trap in white and the background in black.

#### Empty trap finding

The empty traps, designed to block entry of cells, need to be identified so that they can be subtracted from all the other cell traps to subtract background and thereby improve cell pole detection (Fig. S3. D). The empty traps are bright and have a uniform intensity profile. Therefore, to identify them, all cell traps are sorted based on the summed intensity of the pixels in the middle of each cell trap along the y-axis. Of the four brightest in each imaging position, the one with minimal summed derivative, i.e. the most uniform cell trap, is selected. A single best (the brightest and most uniform) empty trap is detected in the first frame of each imaging position. The selection process is repeated one more time among those selected cell traps from different positions to select the top three cell traps of all positions. A virtual empty trap produced by averaging these top three empty traps is used as a global empty trap during background subtraction.

#### Background removal and pole detection

For each cell trap in each frame of each imaging position, the empty trap is subtracted and the intensity is rescaled to increase the intensity resolution of the cell trap region (Fig. S3. E-I to III). The pixel values for each cell trap are summed along the x-axis to provide the intensity profile along the y-axis of the cell trap. The intensity profile is smoothed to flatten noise and its approximate derivative is computed to indicate the locations of the biggest intensity changes. The absolute value of the approximate derivative is computed and smoothed; this provides the profile of the intensity changes and therefore the poles of cells in the cell trap (Fig. S3. E-IV). The peaks of this profile are extracted and the one with the highest y-coordinate is selected as being the foremost cell pole of the cell trap as the back end of the cell trap that is closest to the back channel is the one with the lowest y-coordinate while the front end has the highest y-coordinate.

**Fig. S3.**
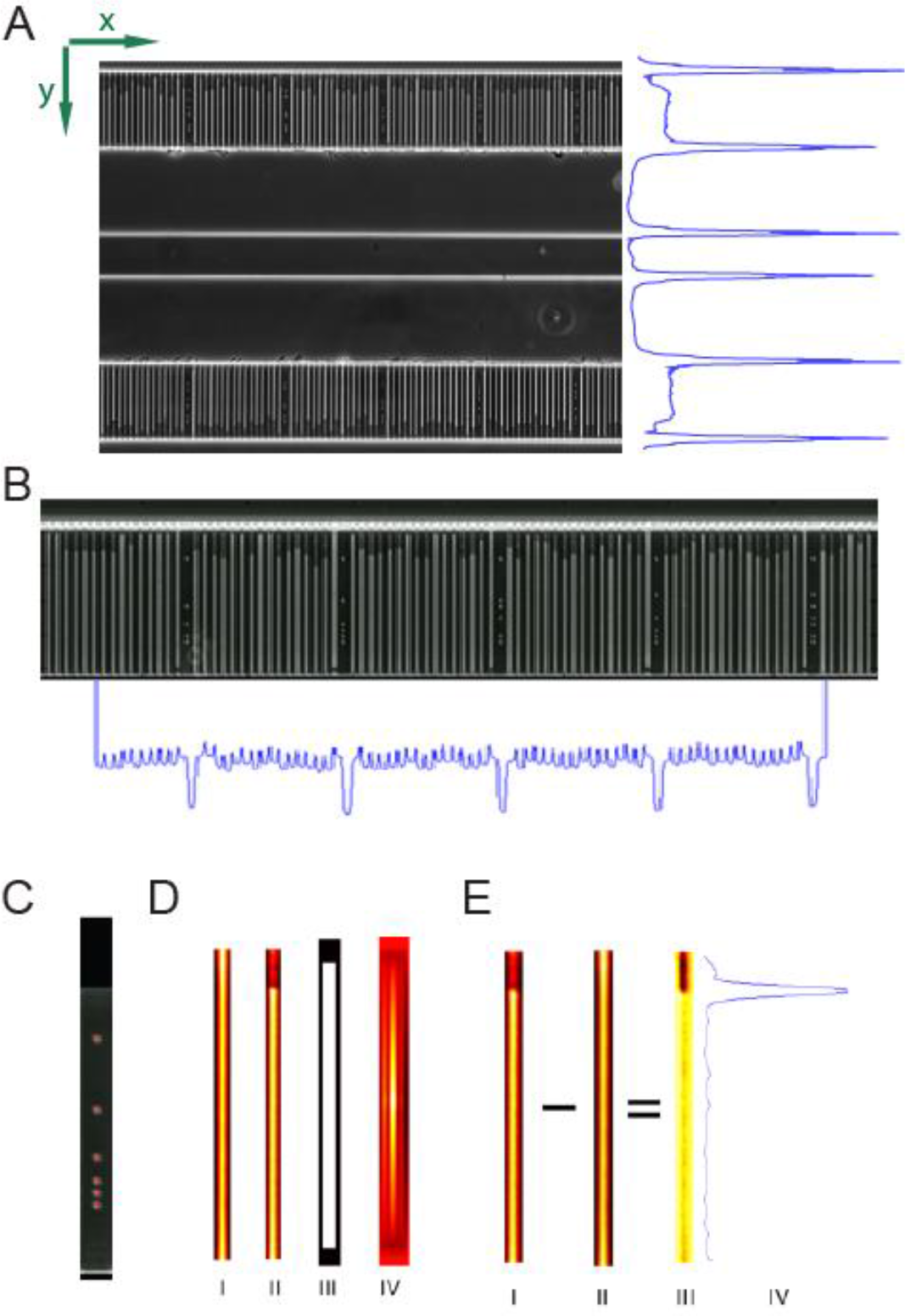
Image processing steps: (A) ROI detection, (B) cell trap detection, (C) precise ROI detection using dot-barcode, (D) empty trap detection, (E) background removal and cell pole detection.

### Data Analysis

Data analysis consists of pole tracking, growth rate calculation with a sliding window, filtering the data for erroneous pole detections and calculation of separation time.

#### Pole tracking

The cell poles are considered to be particles and tracked using uTrack *(32)*.

#### Growth rate calculation with sliding window

The tracking algorithm outputs trajectories each of which contains the location (y-coordinate) of the tracked cell pole in each frame it is tracked. Y-coordinate value also corresponds to the accumulated length of the cell(s) within the particular channel. For exponentially growing cells, the logarithm of cell length has a linear dependence on time. Thus, the growth rate is calculated by fitting a linear regression model to the logarithm of length and the frame number to find the best linear unbiased estimator (BLUE) (i.e. among all unbiased estimators that are linear functions of Y, it has the smallest variance) *(34)*. To be able to detect the changes in growth rate, we estimate the instantaneous growth rate using a sliding window of 20 frames. However, this poses a problem at the beginning of the experiment until the 20^th^ frame. First estimate is from the 5^th^ frame, when for the first time there are enough data points to have a reliable fit. Then, until the 20^th^, the estimate is based on all available frames.

### Data Filtering criteria

Minimum number of data points in a trajectory: If the cell pole detection is not consistently detecting the same cell pole this is likely to result in short trajectories. For this reason, trajectories are discarded if they are shorter than 15 frames.

Maximum filling of the trap: In some experiments, some of the traps might be overfilled during the loading. If a cell trap is loaded more than a certain percent from the beginning, we therefore discard the trajectory from that trap. This also filters the dirt particles close to the open end of the trap that are incorrectly considered to be cell poles. For a trap length of 185 pixels, the trajectory is discarded if the initial y value of the trajectory is > 160.

Minimum growth rate over the entire trajectory: Sometimes the detected cell poles are actually not real cell poles but debris or dead cells loaded in the cell trap, a glare due to dirt on the coverslip or on the illumination path creating an intensity variation that is detected as a cell pole. If the detected cell pole does not move during the experiment, the growth rate calculated from the entire trajectory will be close to zero. Thus, by requiring at least some growth throughout the experiment these false cell poles are filtered out.

### Antibiotics

We have purchased the antibiotics and additives in powder form (Sigma Aldrich, product number is given below as P#) and made aliquots of 500x stock solutions. All the stock aliquots are kept in dark at −80C, and for each experiment, one aliquot is thawed only once, used and excess is discarded to avoid repeating freeze thaw cycles. Different antibiotics requires different solvents. For this reason we dissolved Amoxycillin (P#31586-250MG) and Clavulanate (P#C9874-1G) in phosphate buffer (0.1Mm pH6); Doripenem (P#32138-25MG) in physiological saline (0.85%); Ampicillin, Ciprofloxacin (P#33434-100MG-R), Fosfomycin (P#34089-100MG) and Mecillinam (P#33447-100MG) in deionized water; Trimethoprim (P#46984-250MG), Sulfamethoxazole (P#S7507-10G) and Nitrofurantoin (P#46502-250MG) in DMSO; and Levofloxacin (P#28266-1G-F) initially dissolved in 200mM NaOH and then diluted 2 fold with deionized water. When using Fosfomycin, Glucose-6-Phosphate (P#G7879-500MG), dissolved in deionized water is added to the media to the final 25µg/mL concentration as recommended.

### fASTest Results of Low Cell Density Cultures

To show that fASTest can be used on the low cell density cultures, we have randomly subsampled the original data in one of the experiments, where the wild type *E. coli* is tested with ciprofloxacin (1µg/ml). We have compared subsampling to 951 cells (Fig. S4. A), against subsampling to 85 cells (Fig. S4. B). The separation time increased from 3 minutes to 10 minutes. Most of the experiments in this manuscript were performed with >1000 cells.

**Fig. S4.**
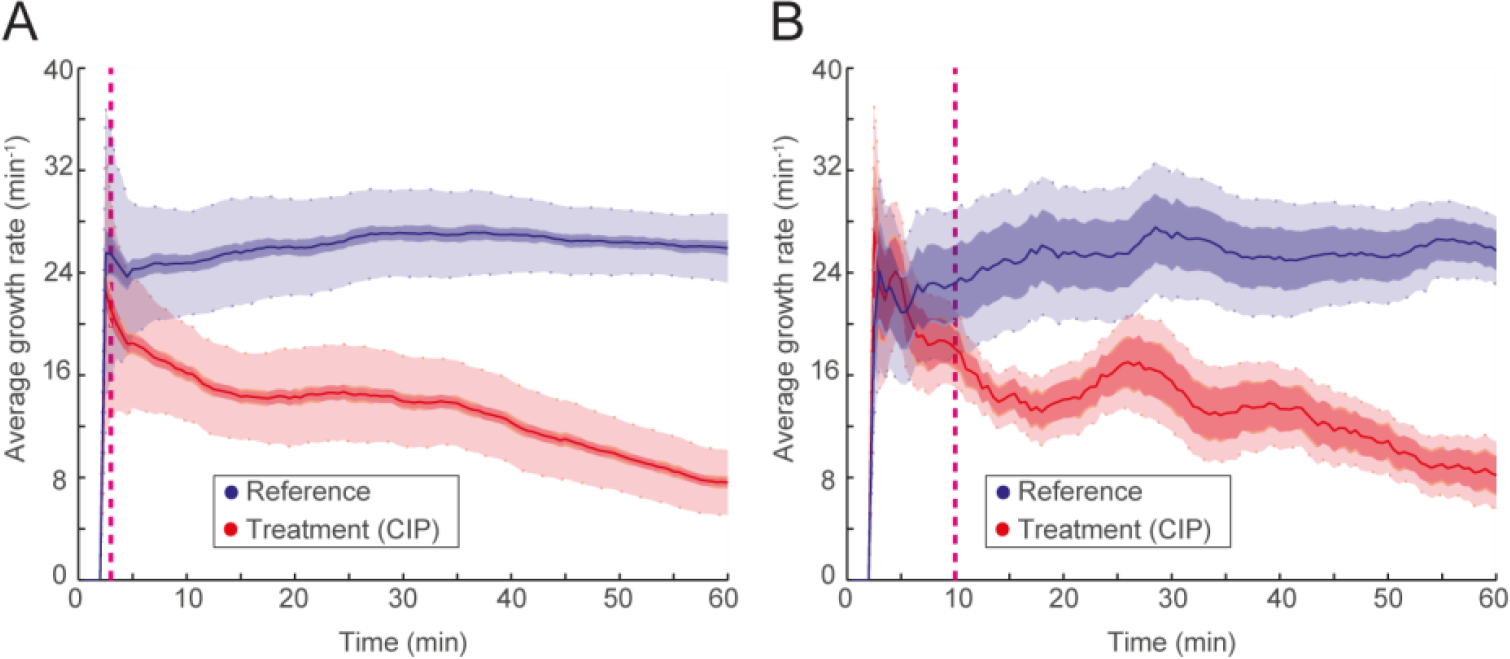
(A) fASTest using 951 cells (reference:409, treatment:502) and (B) 85 cells (reference 44, treatment:41) number of cells is compared.

### fASTest for repeatable detection of drug response using two different antibiotics

To show that the response curves in fASTest are highly repeatable and shape is mainly dependent on the drug being used, we tested susceptible *E. coli* (MG1655) for two different antibiotics and repeated the test for three biological replicates for each antibiotic (Fig. S5).

**Fig. S5.**
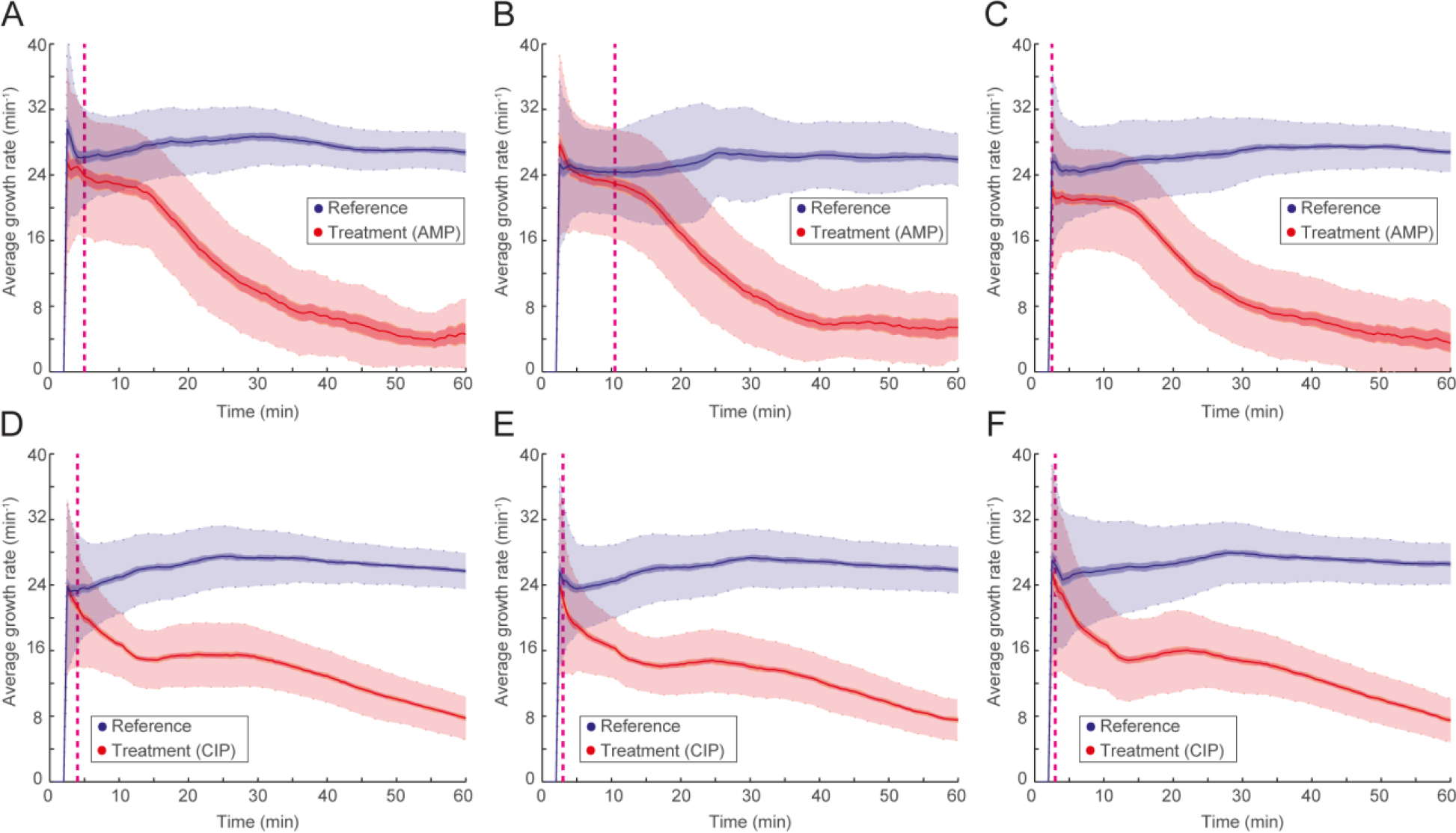
(A, B, C) fASTest of ampicillin and (D, E, F) ciprofloxacin for susceptible *E. coli* (MG1655) biological triplicates.

### fASTest for repeatable detection of drug response in standard laboratory medium and urine

We also tested ciprofloxacin resistant *E. coli* in fASTest using CIP and AMP without reference population to show response to AMP does not differ between the susceptible *E. coli* and CIP resistant strains in three biological triplicates (Fig. S6. A-C). When urine was used as a growth- and test medium instead of Mueller Hinton Broth, the ciprofloxacin resistant *E. coli* showed a very repeatable response to CIP and AMP among three biological replicates (Fig. S6. D-F).

**Fig. S6.**
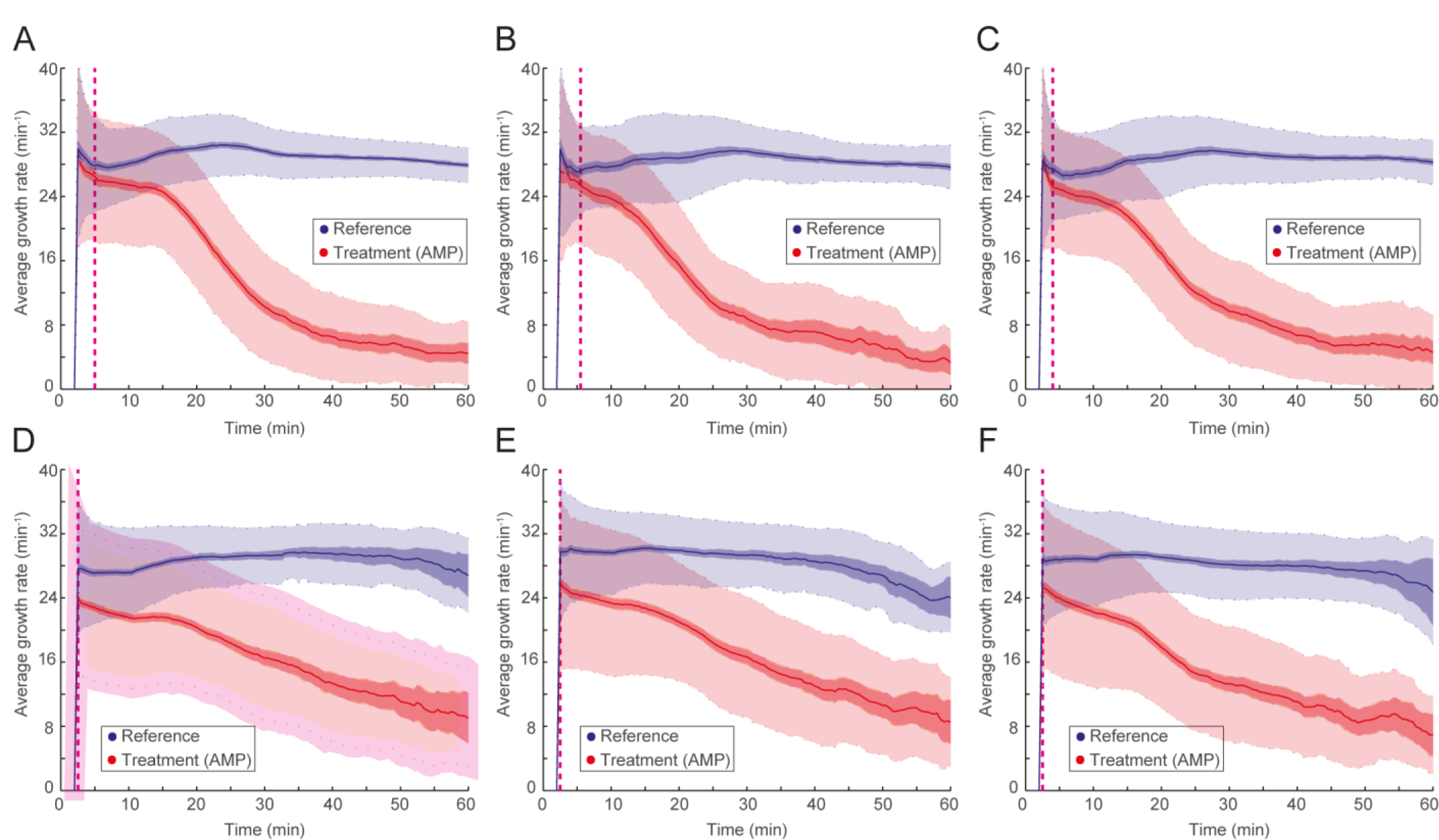
(A-C) fASTest in standard laboratory medium with three biological replicates of CIP resistant strain using CIP (instead of reference) and AMP, and (D-F) fASTest in urine with three biological replicates of CIP resistant strain to CIP and AMP.

### fASTest for different bacterial species: Klebsiella pneumoniae, Staphylococcus saprophyticus

To show that the fASTest can be also be used with other species than *E. coli*, we tested two other species that are the most common after *E. coli* in uncomplicated urinary tract infections: *K. pneumoniae* and *S. saprophyticus* for CIP susceptibility (Fig. S7). *K. pneumoniae* (S.Movie 3), a rod shape gram-negative bacterium grows very similar to *E. coli* (S.Movie 2) in the chip. Also *S. saprophyticus* (S.Movie 4), a bundle forming coccal bacterium, managed to grow even though the division in alternating axes was restricted in one of the axes when restricted into our cell traps. The slower growth rate of *S. saprophyticus* makes the assay both slower and nosier than that observed for the fast-growing *E. coli* and *K. pneumoniae*.

**Fig. S7.**
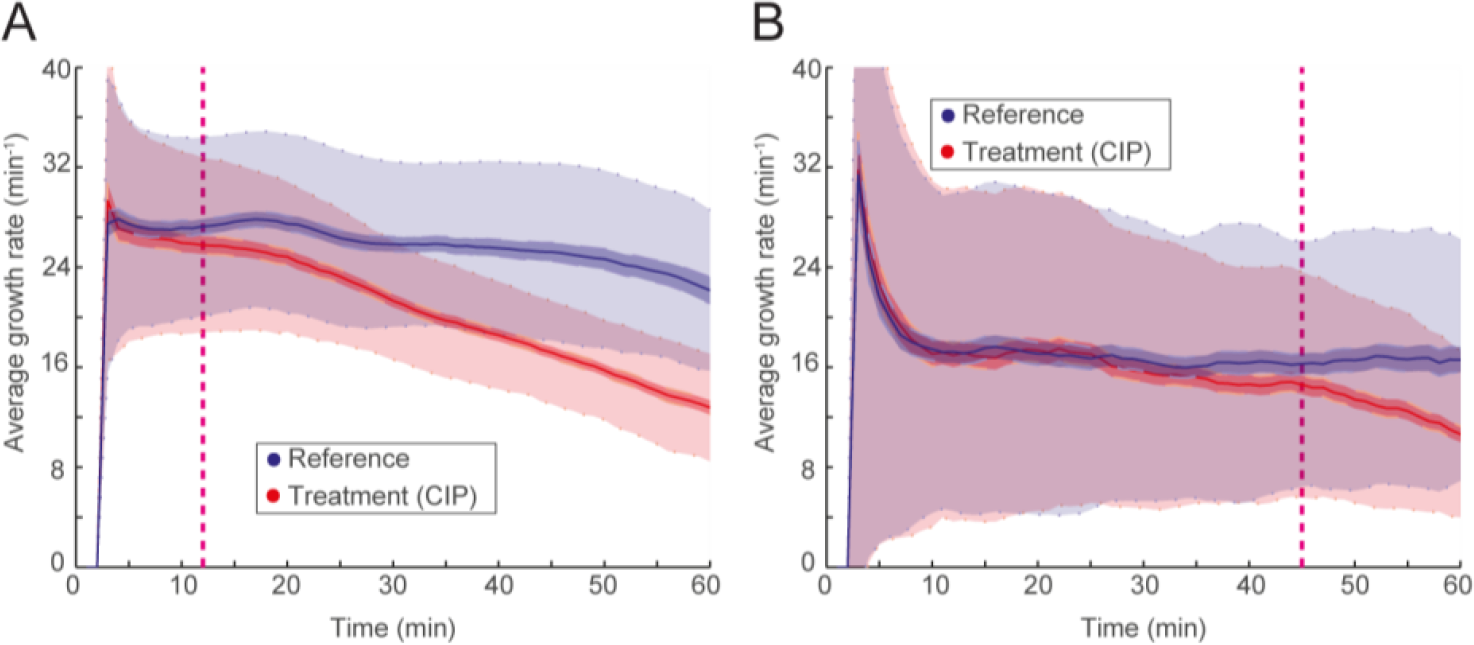
(A) fASTest for *K. pneumoniae* and (B) *S. saprophyticus* for CIP susceptibility detection.

**Movie S1.** Time lapse images of an example FASTest: A time-lapse movie of a FASTest where we loaded the susceptible *E. coli* (MG1655) into the 2GMM and supplied growth media without antibiotic to the top row and growth media with ciprofloxacin (1µg/mL) to the bottom row. This movie is generated from a total of 120 phase contrast images, that are taken with 30 seconds interval. The playback rate is 10 frame per second, i.e. each second in the playback corresponds to 5 minutes in the experiment.

**Movie S2.** Time-lapse imaging (with 100x objective) of susceptible *K. pneumoniae* growing in the microfluidic chip in standard media. This movie is generated from phase contrast images taken with 60sec interval and playback is 300x the speed of the imaging, i.e. each second in the playback corresponds to 300sec in the experiment.

**Movie S3.** Time-lapse imaging (with 100x objective) of susceptible *S. saprophyticus* growing in the microfluidic chip in standard media. This movie is generated from phase contrast images taken with 5sec interval and playback is 100x the speed of the imaging, i.e. each second in the playback corresponds to 500sec in the experiment.

## References and Notes

1. J. J. Kerremans, P. Verboom, T. Stijnen, L. Hakkaart-van Roijen, W. Goessens, H. A. Verbrugh, M. C. Vos, Rapid identification and antimicrobial susceptibility testing reduce antibiotic use and accelerate pathogen-directed antibiotic use, J. Antimicrob. Chemother. 61, 428–435 (2008).

2. G. V. Doern, D. R. Scott, A. L. Rashad, Clinical Impact of Rapid Antimicrobial Susceptibility Testing of Blood Culture Isolates, Antimicrob. Agents Chemother. 21, 1023–1024 (1982).

3. A. Van Belkum, G. Durand, M. Peyret, S. Chatellier, G. Zambardi, J. Schrenzel, D. Shortridge, A. Engelhardt, W. M. Dunne, Rapid clinical bacteriology and its future impact Ann. Lab. Med. 33, 14–27 (2013).

4. S. G. Jenkins, A. N. Schuetz, in Mayo Clinic Proceedings, (2012), vol. 87, pp. 290–308.

5. E. Matuschek, D. F. J. Brown, G. Kahlmeter, Development of the EUCAST disk diffusion antimicrobial susceptibility testing method and its implementation in routine microbiology laboratories, Clin. Microbiol. Infect. 20 (2014), doi: 10.1111/1469-0691.12373.

6. H. Frickmann, W. O. Masanta, A. E. Zautner, Emerging rapid resistance testing methods for clinical microbiology laboratories and their potential impact on patient management Biomed Res. Int. 2014 (2014), doi:10.1155/2014/375681.

7. A. Reece, B. Xia, Z. Jiang, B. Noren, R. McBride, J. Oakey, Microfluidic techniques for high throughput single cell analysis., Curr. Opin. Biotechnol. 40, 90–96 (2016).

8. E. K. Sackmann, A. L. Fulton, D. J. Beebe, The present and future role of microfluidics in biomedical research, Nature 507, 181–189 (2014).

9. C. Murray, O. Adeyiga, K. Owsley, D. Di Carlo, Research highlights: microfluidic analysis of antimicrobial susceptibility., Lab Chip 15, 1226–9 (2015).

10. S. Kim, S. Cestellos-Blanco, K. Inoue, R. Zare, Miniaturized Antimicrobial Susceptibility Test by Combining Concentration Gradient Generation and Rapid Cell Culturing, Antibiotics 4, 455–466 (2015).

11. J. Choi, Y.-G. Jung, J. Kim, S. Kim, Y. Jung, H. Na, S. Kwon, Rapid antibiotic susceptibility testing by tracking single cell growth in a microfluidic agarose channel system., Lab Chip 13, 280–7 (2013).

12. Z. Hou, Y. An, K. Hjort, K. Hjort, L. Sandegren, Z. Wu, Time lapse investigation of antibiotic susceptibility using a microfluidic linear gradient 3D culture device., Lab Chip 14, 3409–18 (2014).

13. J. Choi, J. Yoo, M. Lee, E.-G. Kim, J. S. Lee, S. Lee, S. Joo, S. H. Song, E.-C. Kim, J. C. Lee, H. C. Kim, Y.-G. Jung, S. Kwon, A rapid antimicrobial susceptibility test based on single-cell morphological analysis., Sci. Transl. Med. 6, 267ra174 (2014).

14. D. T. Quach, G. Sakoulas, V. Nizet, J. Pogliano, K. Pogliano, Bacterial Cytological Profiling (BCP) as a Rapid and Accurate Antimicrobial Susceptibility Testing Method for Staphylococcus aureus, EBioMedicine, 1–9 (2016).

15. J. D. Besant, E. H. Sargent, S. O. Kelley, Rapid electrochemical phenotypic profiling of antibiotic-resistant bacteria, Lab Chip 15, 2799–2807 (2015).

16. P. Wang, L. Robert, J. Pelletier, W. L. Dang, F. Taddei, A. Wright, S. Jun, Robust growth of Escherichia coli., Curr. Biol. 20, 1099–103 (2010).

17. C. Slekovec, J. Leroy, N. Vernaz-Hegi, J. P. Faller, D. Sekri, B. Hoen, D. Talon, X. Bertrand, Impact of a region wide antimicrobial stewardship guideline on urinary tract infection prescription patterns, Int. J. Clin. Pharm. 34, 325–329 (2012).

18. G. Schmiemann, E. Kniehl, K. Gebhardt, M. M. Matejczyk, E. Hummers-Pradier, The diagnosis of urinary tract infection: a systematic review., Dtsch. Ä rzteblatt Int. 107, 361–7 (2010).

19. L. L. Marcusson, N. Frimodt-Møller, D. Hughes, Interplay in the selection of fluoroquinolone resistance and bacterial fitness, PLoS Pathog. 5 (2009), doi:10.1371/journal.ppat.1000541.

1. M. Wallden, D. Fange, E. Gregorsson Lundius, Ö. Baltekin, J. Elf, The synchronization of replication and division cycles in individual E. coli cells (in press), Cell (2016).

21. K. Jaqaman, D. Loerke, M. Mettlen, H. Kuwata, S. Grinstein, S. L. Schmid, G. Danuser, Robust single-particle tracking in live-cell time-lapse sequences., Nat. Methods 5, 695–702 (2008).

